## Supplementary Fig. S2 for "Sex-dimorphic expression of extracellular matrix genes in mouse bone marrow neutrophils"

### Figure S2

A WGCNA module cluster dendrogram

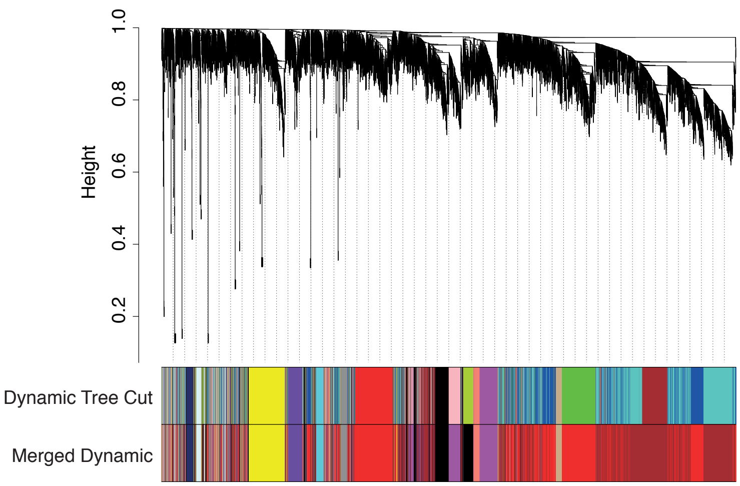


B Heatmap of WGCNA Salmon module gene expression (318 genes; bulk)

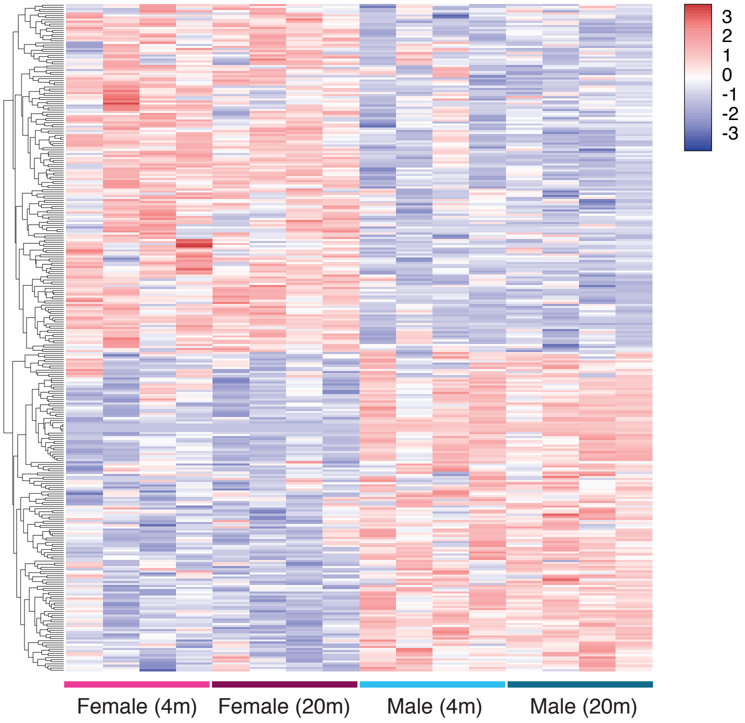
