## Supplementary Fig. S1 for "Sex-dimorphic expression of extracellular matrix genes in mouse bone marrow neutrophils"

### Figure S1

A Enrichment analysis for ECM-related genes in **neutrophil** female-biased genes by RNA-seq

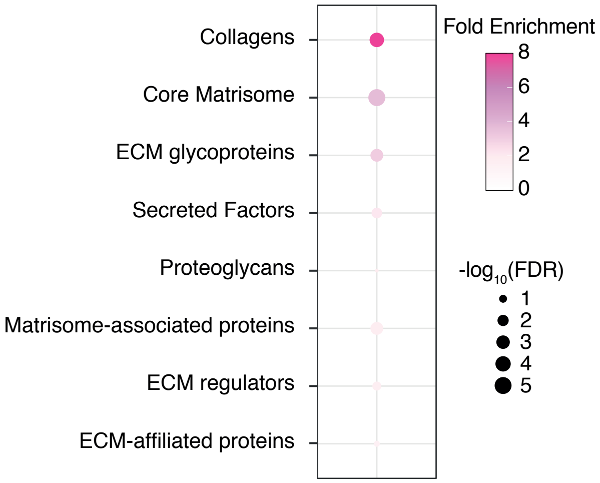


B Ucell analysis of ECM-related genes in **neutrophils** as a function of maturation stage (single cell RNA-seq)

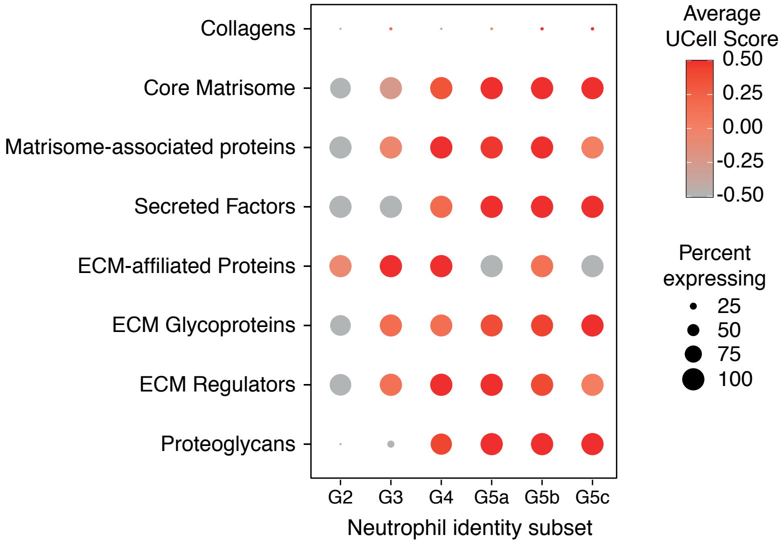


C MDS of **serum** proteomes

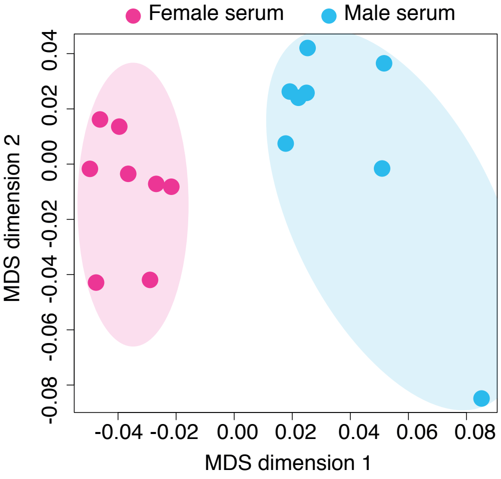


D Heatmap of sex-biased **serum** proteins

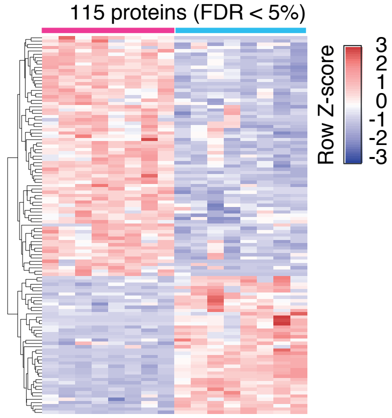


E Enrichment analysis for ECM-related proteins in **serum** female-biased proteins

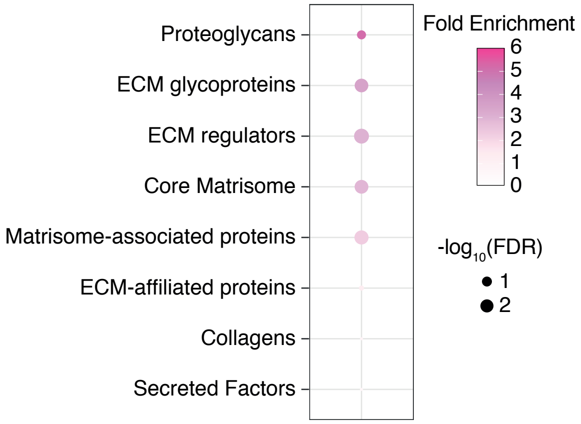
